# Using light and X-ray scattering to untangle complex neuronal orientations and validate diffusion MRI

**DOI:** 10.1101/2022.10.04.509781

**Authors:** Miriam Menzel, David Gräßel, Ivan Rajkovic, Michael Zeineh, Marios Georgiadis

## Abstract

Disentangling human brain connectivity requires an accurate description of neuronal trajectories. However, a detailed mapping of axonal orientations is challenging because axons can cross one another on a micrometer scale. Diffusion magnetic resonance imaging (dMRI) can be used to infer neuronal connectivity because it is sensitive to axonal alignment, but it has limited resolution and specificity. Scattered Light Imaging (SLI) and small-angle X-ray scattering (SAXS) reveal neuronal orientations with microscopic resolution and high specificity, respectively. Here, we combine both techniques to achieve a cross-validated framework for imaging neuronal orientations, with comparison to dMRI. We evaluate brain regions that include unidirectional and crossing fiber tracts in human and vervet monkey brains. We find that SLI, SAXS, and dMRI all agree regarding major fiber pathways. SLI and SAXS further quantitatively agree regarding fiber crossings, while dMRI overestimates the amount of crossing fibers. In SLI, we find a reduction of peak distance with increasing out-of-plane fiber angles, confirming theoretical predictions, validated against both SAXS and dMRI. The combination of scattered light and X-ray imaging can provide quantitative micrometer 3D fiber orientations with high resolution and specificity, enabling detailed investigations of complex tract architecture in the animal and human brain.

## Introduction

Unraveling the complex nerve fiber network in the brain is key to understanding its function and alterations in neurological diseases. The detailed reconstruction of multiple crossing, long-range nerve fiber pathways in densely-packed white matter regions poses a particular challenge. *Diffusion magnetic resonance imaging* (*dMRI*) is currently used to derive neuronal orientations *in vivo*. However, with voxel sizes typically down to a few hundred micrometers in post-mortem human brains (***Calabrese et al., 2018**; **Roebroeck et al., 2018***), the resolution is insufficient to resolve individual nerve fibers, and the signal is affected by all brain structures, not only axons. Moreover, the possibly hundreds of fibers within a voxel might have complicated geometries, e.g. crossing or kissing fibers, which poses a particular challenge. Especially notable is that structural connectivity and wiring diagrams of the brain, obtained from dMRI measurements and subsequent fiber tractography, contain a large percentage of false-positive fiber tracts (***Maier-Hein et al., 2017; Schilling et al., 2019; Maffei et al., 2022***), indicating a poor specificity in detecting actual fiber tracts.

*Small-angle X-ray scattering* (*SAXS*) provides myelinated nerve fiber orientations by studying the anisotropy of myelin diffraction (Bragg) peaks in X-ray scattering patterns (***Figure 1A,C*** – ***Georgiadis et al., 2020**; **Georgiadis et al., 2021***). These are generated by the interaction of the incoming X-ray photons with the layered structure of the myelin sheath, which surrounds nerve fibers in the white matter. The method can be tomographic (SAXS tensor tomography – ***Liebi et al, 2015**; **Gao et al, 2019; Georgiadis et al, 2021***), and 3D-scanning SAXS (3D-sSAXS) can provide 3D distributions of axon orientations in tissue sections (***Georgiadis et al, 2020***). Recent studies further revealed that SAXS can exploit the modulations in the azimuthal position of the myelin-specific Bragg peaks to resolve crossing nerve fiber populations across species (***Georgiadis et al., 2022***).

**Figure 1.**
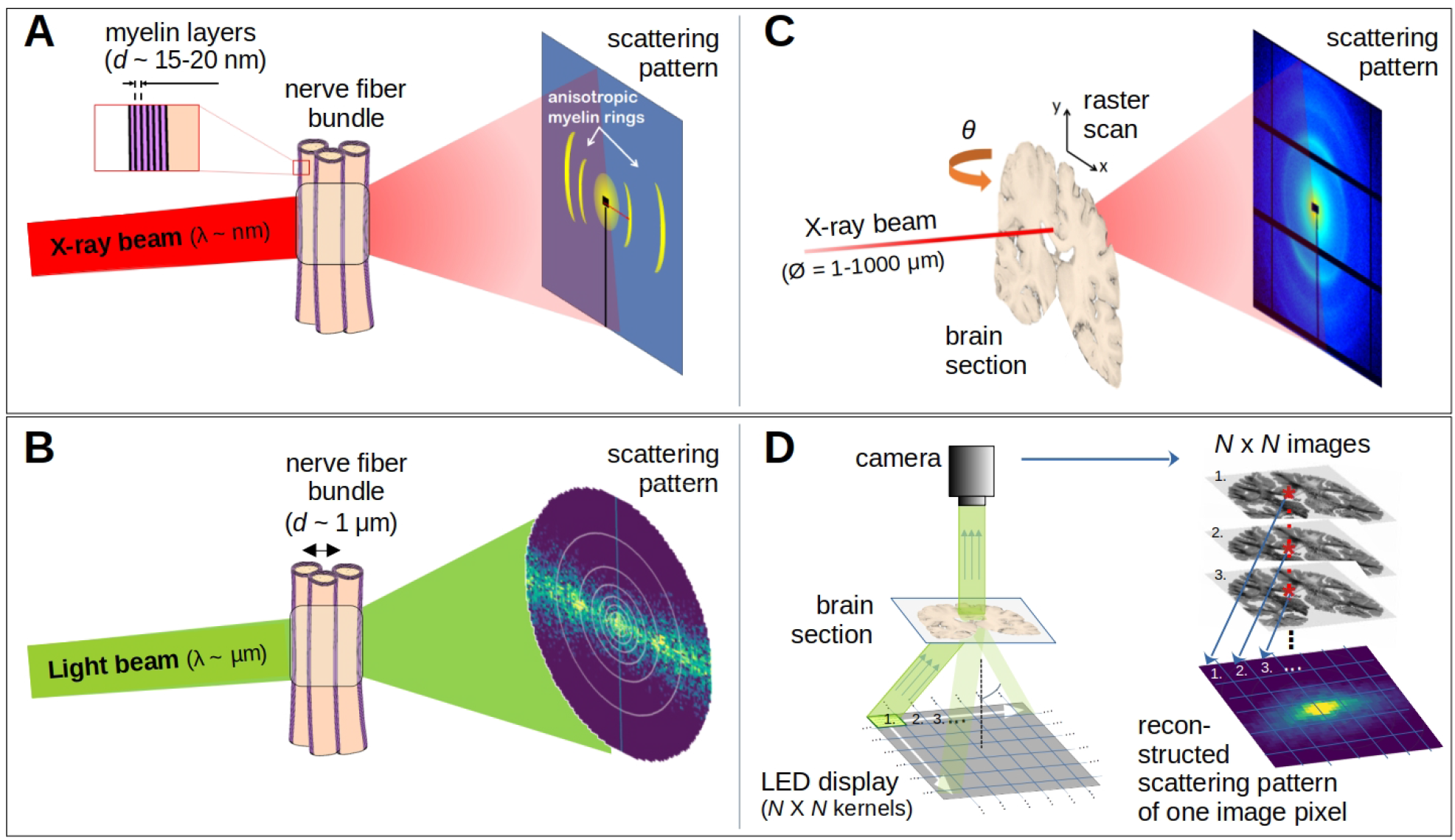
Comparison of X-ray scattering (top) and light scattering (bottom) for analyzing nerve fiber structures. (**A**) Principle of X-ray scattering on a nerve fiber bundle, whereby myelin’s periodicity results in a predictable ring, strongest perpendicular to the in-plane fiber orientation. (**B**) Principle of light scattering on a nerve fiber bundle, which similarly yields scattered photons perpendicular to the in-plane fiber orientation. (**C**) Schematic drawing of a 3D-scanning SAXS measurement of a brain section, in which raster scanning from multiple angles reconstructs 3D fiber orientation distributions in each point of illumination. (**D**) Schematic drawing of an SLI scatterometry measurement of a whole brain section (left) and the reconstruction of a scattering pattern shown for one selected image pixel (right), which can be done over the entire image simultaneously.

The scattering of visible light can also be used to reveal crossing nerve fiber orientations (***Figure 1B – Menzel et al., 2020a,b***) as it is sensitive to directional arrangements of neuronal axons (~μm diameter). In *Scattered Light Imaging (SLI) (**Menzel et al., 2021a,b**; **Reuter and Menzel, 2020***) the sample (brain section) is illuminated from many different angles and a camera captures an image of the brain section (***Figure 1D*** left), in which the intensities of each image pixel vary with the angle of illumination. In this way, a scattering pattern is generated for each micron-sized image pixel (***Figure 1D*** right). SLI has been shown to reliably reconstruct up to three in-plane fiber orientations for each image pixel (with an accuracy of +/-2.4°; ***Menzel et al., 2021a***).

Hence, a combination of 3D-sSAXS and SLI, with the high specificity to myelinated fibers of the former, and the high-resolution capabilities of the latter, can serve as gold standard for imaging complex nerve fiber orientations in the brain with micrometer resolution.

Here, we present combined 3D-sSAXS and SLI measurements on the same tissue samples (coronal sections from vervet monkey and human brains) and compare them to dMRI outcomes. To capture multiple possible fiber scenarios, we examine brain regions with both unidirectional and complex/crossing fibers – the corpus callosum and corona radiata, respectively. Evaluation of combined 3D-sSAXS and SLI in a vervet brain section provides a unique cross-validation, but also a very detailed mapping of the single and crossing fiber orientations. Comparison of the results on the human brain sample enables validation of dMRI-derived orientations, which offers the possibility of *in vivo* translation. An overestimation of the number of fiber crossings is identified in dMRI. Furthermore, we enhance the interpretation of out-of-plane fibers in SLI, using the 3D-fiber orientations from SAXS and dMRI as reference. The presented framework can be used to provide reliable axonal orientations, validate dMRI results, and deliver more accurate brain connectivity maps of the animal and human brain.

## Results

### Light and X-ray scattering patterns are specific to different fiber configurations

To better understand how light and X-ray scattering patterns correspond to each other for different nerve fiber configurations, we analyzed the scattering patterns from SLI and SAXS measurements in a vervet monkey brain section. ***Figure 2*** shows the resulting scattering patterns for four representative points (marked with asterisks in B): (i) unidirectional in-plane fiber bundle in the corpus callosum, (ii) two crossing fiber bundles in the corona radiata, (iii) a sightly through-plane inclined fiber bundle in the fornix, and (iv) a steep out-of-plane fiber bundle in the cingulum. The orientation information is encoded in the variation of the signal intensity as a function of the azimuthal angle *φ* (going in a circle around the pattern, cf. ***Figure 2C***(i)), plotted as *azimuthal profile* under each scattering pattern in ***Figure 2C**. **Figure 2–figure supplement 1*** shows average, maximum, minimum, mean peak prominence and mean peak width of the azimuthal profiles for each pixel measured with SAXS and SLI.

**Figure 2.**
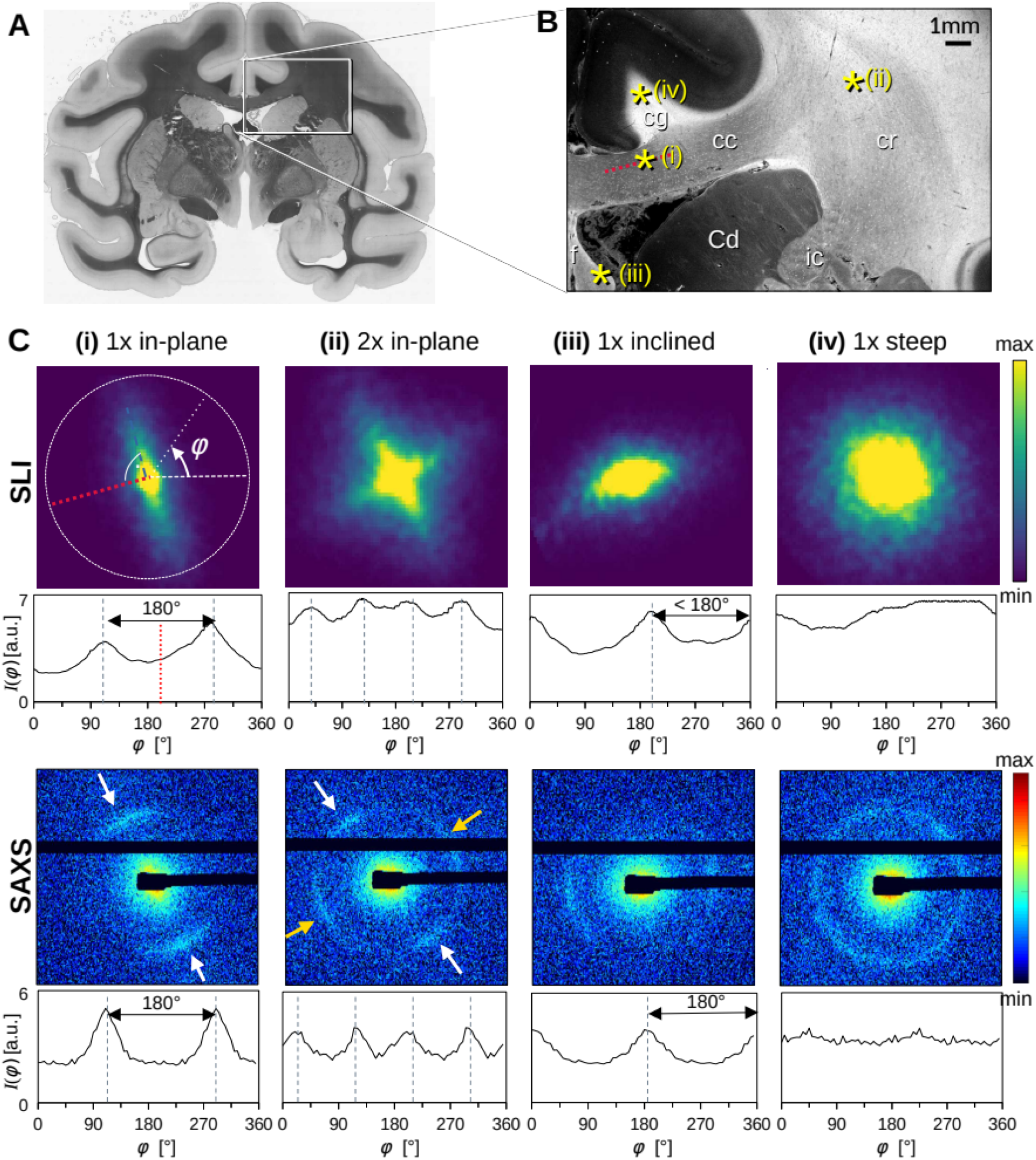
Scattering patterns obtained from SLI scatterometry (px = 3μm) and SAXS (px = 100μm) on a 60μm-thick vervet monkey brain section at a coronal plane between amygdala and hippocampus (no. 511). (**A**) Transmittance image of the whole brain section. (**B**) Average scattered light intensity of the investigated region (cc: corpus callosum, cr: corona radiata, cg: cingulum, Cd: caudate nucleus, f: fornix, ic: internal capsule). Yellow asterisks indicate the points corresponding to the scattering patterns in C. (**C**) SLI and SAXS scattering patterns, with azimuthal profiles plotted beneath each pattern, obtained from the pixels indicated in B. (i) unidirectional in-plane fiber bundle in the corpus callosum, with peaks perpendicular to the fiber orientation in red, lying 180° apart, (ii) two in-plane crossing fiber bundles in the corona radiata, (iii) slightly inclined fiber bundle in the fornix, with SLI peaks <180° apart, and SAXS peaks 180° apart but with lower peak height, (iv) highly inclined fiber bundle in the cingulum. The following figure supplement is available for figure 2: **Figure supplement 1.** Parameter maps obtained from SAXS and SLI azimuthal profiles for vervet monkey brain section no. 511.

While the SLI scattering patterns show contiguous signal intensity (from center out), the strongest SAXS signal (Bragg peaks) appears along the Debye-Scherrer ring (arrows in ***Figure 2C***), at a specific distance (q-value) from the center of the pattern that corresponds to the myelin layer periodicity (here 17.5nm) (***Georgiadis et al., 2021***).

For in-plane nerve fibers, i.e. nerve fibers that mostly lie within the section plane, the strongest signal in both SLI scatterometry and SAXS is perpendicular to the fiber orientation (red dashed lines in ***Figure 2B and C***(i)), shown in the azimuthal profile as peaks separated by 180°. For the two in-plane crossing fiber bundles in the corona radiata (ii), the peaks in the SLI and SAXS azimuthal profiles similarly indicate the fiber orientations, with each bundle producing two peaks separated by 180° (white/yellow arrows).

For partly out-of-plane fibers, i.e. fibers that have a certain angle with respect to the section plane, such as those in the fornix, the peaks in the SAXS azimuthal profiles are still 180° apart - owing to the center-symmetry of the pattern -, but become less pronounced with increasing out-of-plane fiber angle (compare peak height of SAXS, ***Figure 2C***(i) vs. (iii)). In contrast, the between-peak distance in the SLI azimuthal profiles decreases with increasing fiber inclination (SLI, ***Figure 2C***(iii)), as also predicted by simulation studies (***Menzel et al., 2020a***). For out-of-plane fibers that run almost perpendicular to the section plane (***Figure 2*** point (iv), cingulum), the SLI scattering pattern becomes almost radially symmetric and SAXS demonstrates a symmetric ring, neither with visible peaks in the azimuthal profile. In such cases, the information about the in-plane fiber orientations is limited, whereas the out-of-plane angle can be determined using 3D-sSAXS (***Georgiadis et al., 2020***), and approximated in SLI (***Menzel et al., 2021***).

### SAXS and SLI resolve crossing fibers and show high inter-method reproducibility

We then sought to more precisely compare the in-plane nerve fiber orientations derived from the peak positions in the SAXS and SLI azimuthal profiles, examining the same ~1×2cm^2^ region of the vervet brain (***Figure 3*** and ***Figure 3–figure supplement 1***). Given the ~33x higher resolution of SLI over SAXS in the presented measurements (3μm vs. 100μm pixels), smaller nerve fiber bundles e.g. in the head of the caudate nucleus (yellow arrow) can be traced. Conversely, out-of-plane nerve fibers in the cingulum (cg), are more sensitively depicted by SAXS.

**Figure 3.**
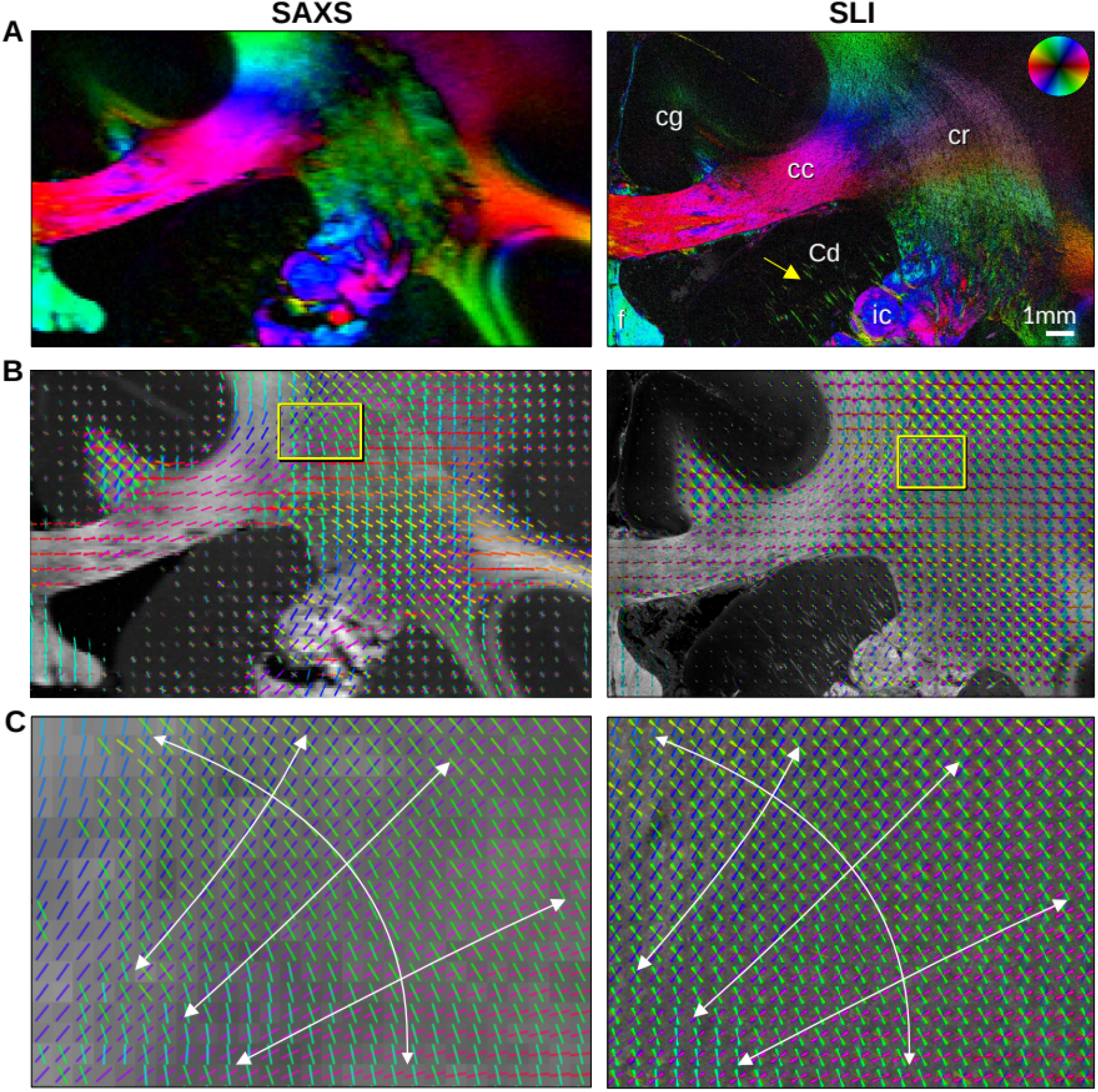
In-plane nerve fiber orientations from SAXS and SLI measurements of vervet monkey brain section no. 511. (**A**) Fiber orientation maps showing the predominant fiber orientation for each image pixel in different colors (see color wheel in upper right corner): px = 100μm (SAXS), px = 3μm (SLI). (cc: corpus callosum, cr: corona radiata, cg: cingulum, Cd: caudate nucleus, f: fornix, ic: internal capsule). (**B**) Fiber orientations displayed as colored lines for 5×5 px (SAXS) and 165×165 px (SLI) superimposed. The length of the lines is weighted by the averaged scattered light intensity in SAXS and SLI, respectively. (**C**) Enlarged region of the corona radiata, showing fiber orientations as colored lines for 1×1 px (SAXS) and 33×33 px (SLI) superimposed. The white arrows indicate the main stream of the computed fiber orientations. The following figure supplement is available for figure 3: **Figure supplement 1.** In-plane fiber orientations from SAXS and SLI measurements of vervet brain section no. 501.

Despite the different resolutions, the in-plane nerve fiber orientations are highly coincident, not only for unidirectional fibers, but also for fiber crossings (colored lines in ***Figure 3B-C**, **Figure 3–figure supplement 1D-E***), where each vector glyph covers orientations from a grid of 165×165 measured pixels that are visually overlaid in SLI, vs. a 5×5 pixel grid in SAXS (***Figure 3B***). Further zooming in shows a concordant fiber course in the highly complex corona radiata architecture (***Figure 3C, Figure 3–figure supplement 1E**):* the fibers of the corpus callosum fan out (blue/magenta) while crossing the ascending internal/external capsule including thalamo-cortical projections (green).

To quantitatively compare the in-plane fiber orientations, the SAXS images were linearly registered onto the SLI images, and pixels in which both techniques yield one or two fiber orientations were compared to each other: For each image pixel, the fiber orientations were subtracted (SLI – SAXS), taking the minimum of the two possible pairings in regions with crossing fibers (***Figure 4***). **Figure 4C** shows the image pixels for which both techniques yield a single fiber orientation (magenta) or two fiber orientations (green). ***Figure 4A*** shows very small angular differences that appear to be uniformly distributed, depicted as absolute angular differences in ***Figure 4B***. While in-plane and slightly inclined fibers (corpus callosum and fornix) as well as major parts of crossing fibers in the corona radiata show mostly differences less than 10°, highly inclined fibers in the cingulum and the corona radiata show absolute differences of 20° and more (white arrows).

**Figure 4.**
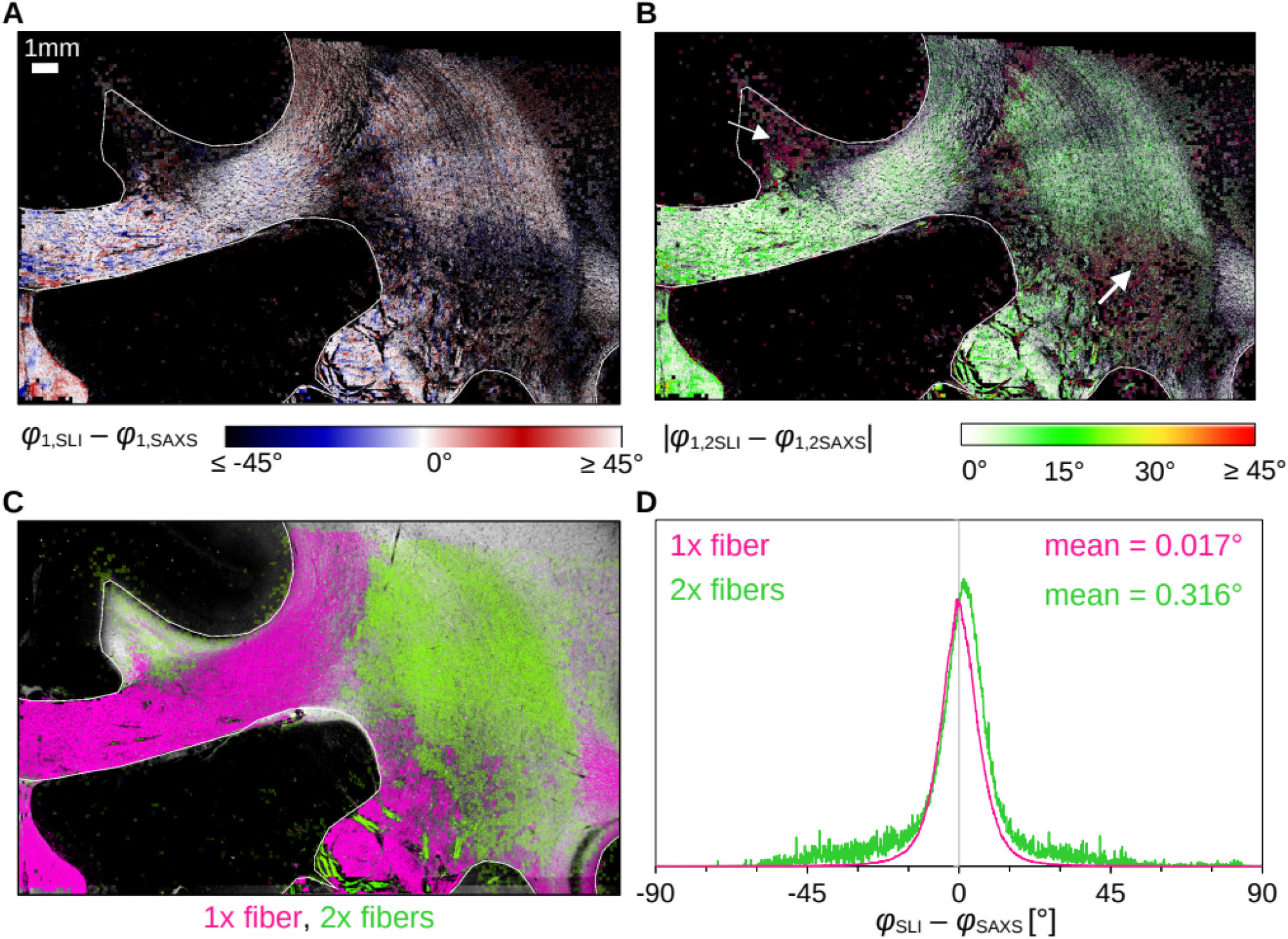
Angular difference between nerve fiber orientations (SLI – SAXS) for vervet monkey brain section no. 511. For evaluation, the SAXS image was registered onto the SLI image and only regions where both techniques yield one or two fiber orientations were considered. (**A**) Angular difference displayed for one of maximum two predominating fiber orientations in each pixel. (**B**) Angular absolute difference displayed for each image pixel. (**C**) Regions with one or two fiber orientations displayed in different colors (magenta=1, green=2 orientations). (**D**) Histograms showing the angular difference for pixels with one and two fiber orientations, evaluated in white matter regions excluding the fornix (see regions delineated by white lines in A-C).

The distribution of angular differences for white matter with one and two fiber orientations is shown in ***Figure 4D*** (histograms in magenta and green, respectively). The two histograms show a distribution around zero degrees (one fiber orientation: mean ~0.017°, median absolute ~4.1°; two fiber orientations: mean ~0.316°, median absolute ~5.6°). While regions with one fiber orientation yield differences between +/-30° maximum, regions with two fiber orientations show multiple outliers with differences of +/-45° and more. As 33×33 SLI pixels with different fiber orientations correspond to one SAXS pixel with a single fiber orientation, larger differences between in-plane fiber orientations are expected, especially in regions with highly varying fiber orientations.

### Diffusion MRI tends to overestimate fiber orientations in the human brain

Next, we aimed to extend our findings to the human brain and to the validation of diffusion MRI (dMRI) fiber orientations. To enable the analysis of regions with both unidirectional and crossing fibers, we selected a ~1cm thick human brain sample that contains parts of the corpus callosum (cc), the cingulum (cg), and the corona radiata (cr) (***Figure 5A***). After high-resolution multi-shell dMRI scanning, we computed traditional diffusion tensors to yield the main fiber orientations (***Figure 5C*** left), and multi-shell multi-tissue constrained spherical deconvolution (***Jeurissen et al., 2014***) to map fiber orientation distribution (***Figure 5D, Figure 5–figure supplement 1)***, including regions with highly aligned fibers as well as distinct fiber crossings.

**Figure 5.**
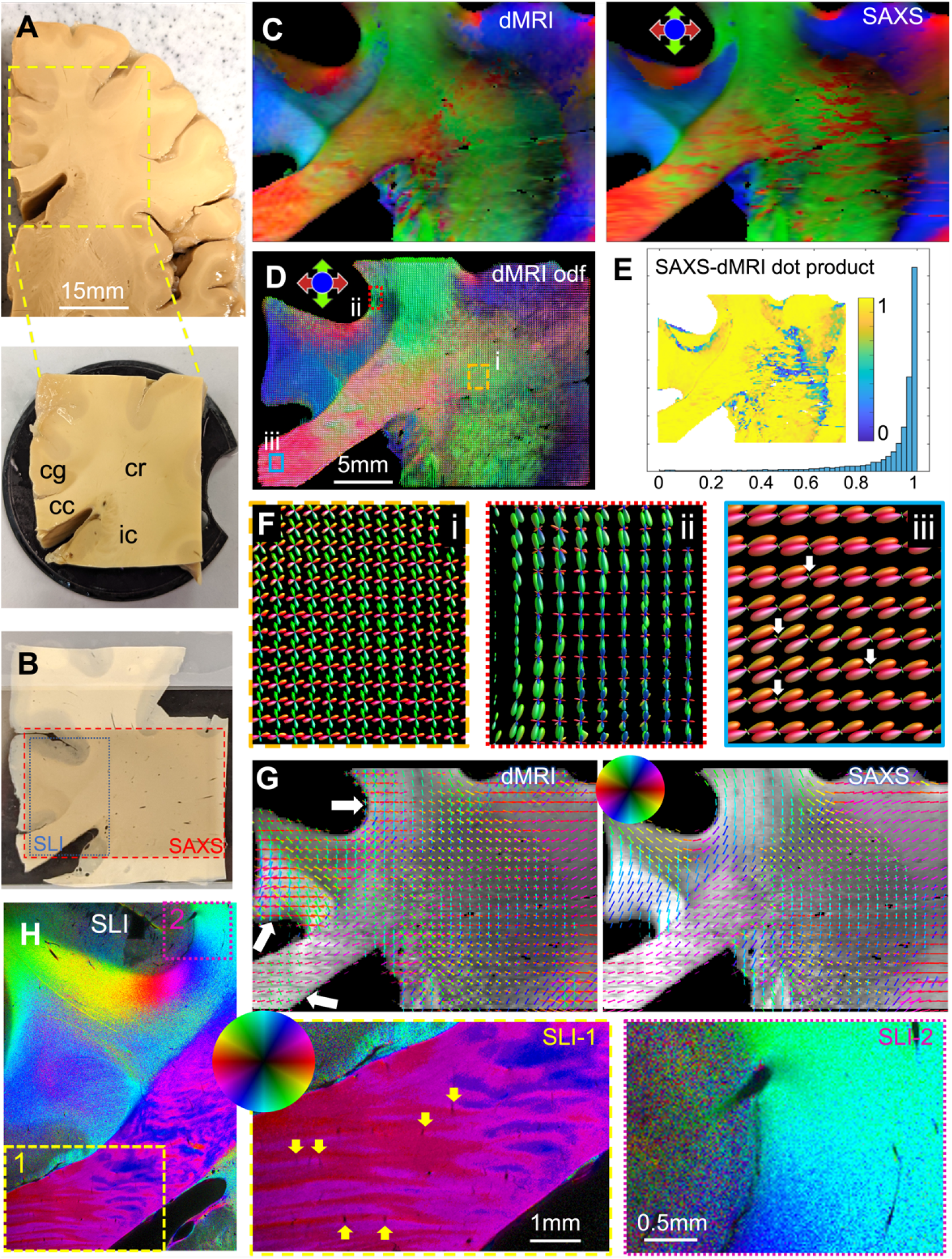
Diffusion MRI measurement of a 3.5×3.5×1cm^3^ human brain specimen (200μm voxel size) in comparison to measurements with 3D-sSAXS (150μm pixel size) and SLI (3μm pixel size) of a 80μm-thick brain section. (**A**) Human brain specimen; the bottom image shows the sample measured with dMRI (cc: corpus callosum, cg: cingulum, cr: corona radiata, ic: internal capsule). (**B**) Posterior brain section with regions measured by 3D-sSAXS (red rectangle) and SLI (blue rectangle). (**C**) Registered main 3D fiber orientations from dMRI (left) and 3D-sSAXS (right) for the brain section. (**D**) Orientation distribution functions from dMRI, with zoomed-in regions surrounded by rectangles shown in (F). (**E**) Vector dot product of the dMRI and 3D-sSAXS main fiber orientations, as histogram and map of the studied area. (**F**) The enlarged regions from (D) show the fiber orientation distributions in the corona radiata (rectangle i, orange), a subcortical U-fiber bundle (rectangle ii, red), and the corpus callosum (rectangle iii, cyan). (**G**) In-plane fiber orientation vectors for dMRI (left) and SAXS (right) superimposed on mean SAXS intensity. Vectors of 5×5 pixels are overlaid to increase visibility. Zoomed-in images of the corona radiata region from both methods are shown in **Figure 5 – Figure supplement 3C.** (**H**) In-plane fiber orientations from SLI (multiple fiber orientations are displayed as multicolored pixels), with zoomed-in areas in boxes (1) and (2); the arrows in box (1) indicate blood vessels. For better readability, fiber orientations in the gray matter are not shown in subfigures C-G. The following figure supplements are available for figure 5: Figure supplement 1-3.

To validate the dMRI-derived fiber orientations, we measured two 80μm-thick vibratome sections (one from the anterior side, ***Figure 5,*** and one from the posterior side, ***Figure 5–figure supplement 2***) with 3Dscanning SAXS and computed fiber orientation distributions (***Georgiadis et al., 2015, 2020***). To enable a quantitative comparison of the 3D fiber orientations obtained from dMRI and 3D-sSAXS, the dMRI sections corresponding to the physical SAXS-scanned sections (cf. ***Figure 5B***, red rectangle) were identified, and linearly registered to the SAXS data sets. The main fiber orientations per pixel for dMRI and 3D-sSAXS (***Figure 5C***) show a high correlation, similar to what has been shown in ***Georgiadis et al. (2020)***, with a dot product approximating unity (***Figure 5E***), and a median angular difference of 14.4° over all voxels (9.1° over voxels with fractional anisotropy (FA) >0.2 for both methods).

We then performed a more detailed analysis including crossing fibers. First, in the challenging region of the corona radiata, where multiple fiber crossings occur, the dMRI orientations seem to be in high agreement with the directly structural X-ray scattering (***Figure 5G*** and ***Figure 5–figure supplement 3B*** left): the two methods have a median angular difference of 5.6° in the primary orientation, and 6.0° in the secondary orientation (overall median angular difference 5.8°). This shows that diffusion MRI has the sensitivity to accurately resolve multiple fiber orientations per voxel, see also ***Figure 5D,F*** (rectangle i, orange). Next, we turned our focus to areas that appear to have relatively homogeneous fiber populations in SAXS, such as the corpus callosum. The main fiber orientations in these regions were again in high agreement between the two methods, with a median angular difference of 5.7° (***Figure 5–figure supplement 3B*** right).

However, there is a striking difference when it comes to resolving secondary orientations. Diffusion MRI seems to also show multiple fiber orientations per voxel, with a secondary fiber population perpendicular to the main one (albeit with much smaller magnitude), in areas where X-ray scattering shows homogeneous fiber orientations, exemplified in the corpus callosum and in the subcortical white matter nearby the cingulate and the callosal sulci (arrows in ***Figure 5G,F***). Referencing these regions in the higher-resolution SLI (px=3μm, ***Figure 5H***), we confirm the X-ray scattering results and do not observe a second fiber population perpendicular to the main one. What can be seen in this micrometer imaging, however, are vessels running perpendicular to the fiber orientations in the corpus callosum area (see yellow arrows in region 1), which might be one of the reasons for the additional fiber directions obtained from dMRI.

We then proceeded to quantify this effect over the entire white matter of the posterior brain section. Comparing the SAXS and dMRI secondary orientations, we observed a 104% (more than double) increase in the voxels with multiple orientations in dMRI. More specifically, secondary fiber orientations within a single voxel were detected in 31% of the total number of voxels by SAXS vs. 64% of the total number of voxels by dMRI. The anterior brain section similarly showed a 40% increase (***Figure 5–figure supplement 2***).

### Experimental validation of out-of-plane fiber orientations in SLI

While SLI determines the in-plane fiber orientation with high precision, out-of-plane fiber orientation (inclination) is challenging. Theory suggests that the fiber inclination is directly related to the distance between the two peaks in the SLI azimuthal profile (cf. upper ***Figure 2C***). The peak distance should decrease with increasing inclination, as indicated by the dashed curves in ***Figure 6G***, which were computed from simulated SLI azimuthal profiles for fiber bundles with different inclinations (***Menzel et al., 2021a,*** Figure 7d). The combined measurement of SLI and 3D-sSAXS enables testing of this prediction, given the very high agreement of 3D-sSAXS and dMRI in the human brain sample in regions of out-of-plane fibers (***Figure 5C-E***).

**Figure 6.**
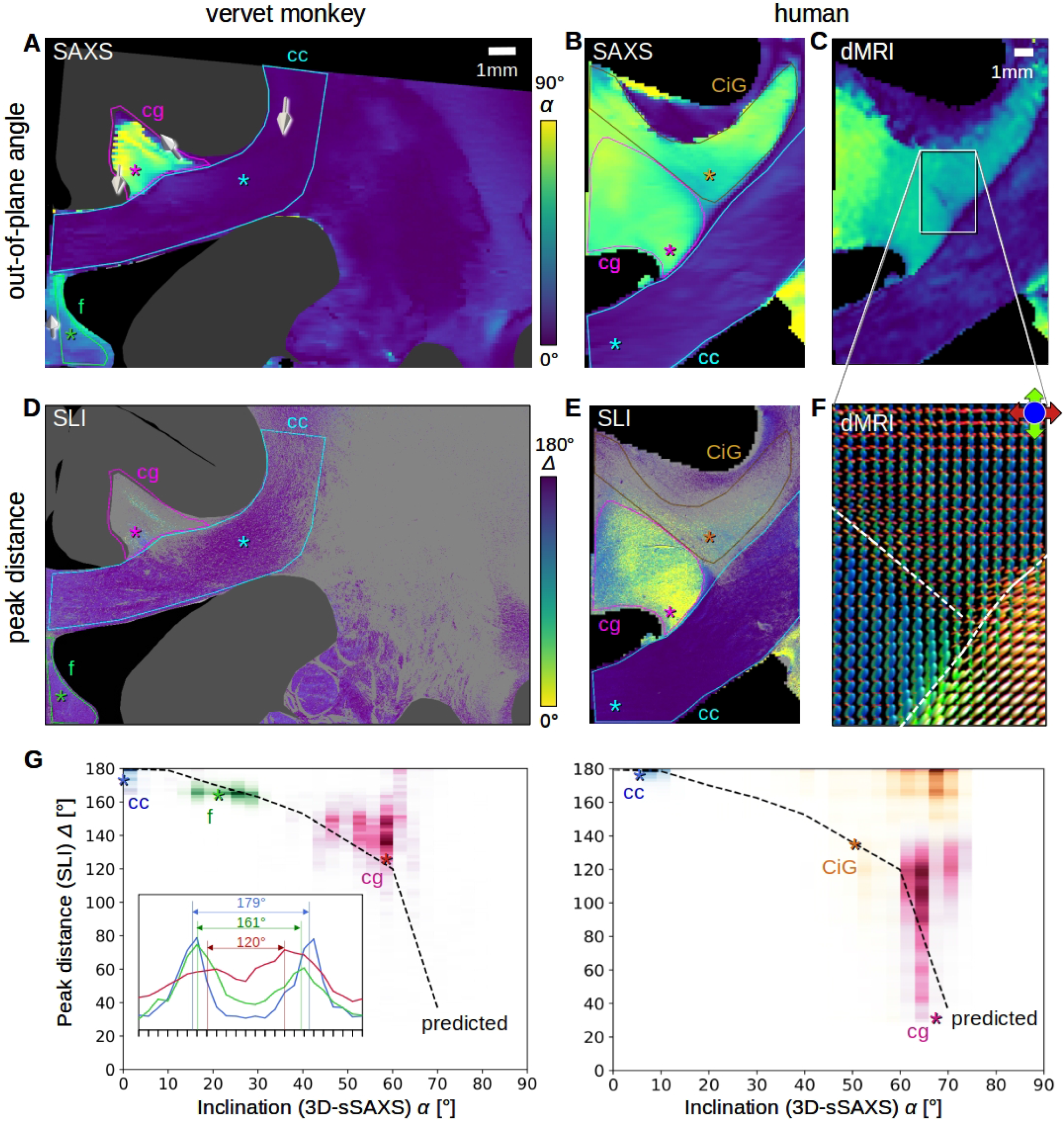
Pixel-wise comparison of 3D-sSAXS/dMRI fiber inclinations and SLI peak distances. The images on the left show the analysis of one vervet brain section (no. 511, cf. **Figure 2B**); the images on the right show the analysis for one human brain section (posterior section, cf. blue rectangle in **Figure 5B**). 3D-sSAXS and dMRI images were registered onto the corresponding SLI images (the fornix in the vervet brain section was additionally shifted between the SLI and 3D-sSAXS images to account for the slight misalignment between the registered images in this region); only regions with unidirectional fibers were evaluated. (**A,B**) 3D-sSAXS fiber inclination angles of the vervet and human brain section shown in different colors for the white matter (blue: in-plane, yellow: out-of-plane, gray: gray matter). The arrows indicate the 3D-orientation of the nerve fibers in four selected regions of the vervet brain section. (**C**) Corresponding dMRI fiber inclination angles of the human brain sample. (**D,E**) Distance between two peaks in the corresponding SLI azimuthal profiles (cf. inset in G). Only profiles with one or two peaks were evaluated (other pixels are shown in gray). Regions used for the pixel-wise comparison with 3D-sSAXS are surrounded by colored outlines; asterisks mark three representative pixels. (**F**) dMRI orientation distribution functions of the region marked in C. The dashed lines indicate separation into the three regions in E (cg – cingulum, CiG – cingulate gyrus, cc - callosum). (**G**) SLI peak distance plotted against the 3D-sSAXS inclination for the corresponding regions marked in D and E (data points are displayed in similar colors as the corresponding outlines). The inset shows the SLI azimuthal profiles for the three representative pixels in the vervet brain section (marked by colored asterisks in D and G). The SLI profiles were centered for better comparison; the ticks on the inset x-axis denote azimuth steps of 15°. The dashed curves indicate the predicted SLI peak distance obtained from simulated scattering patterns of fiber bundles with different inclinations (**Menzel et al., 2021a,** Figure 7d).

We performed a pixel-wise comparison of the out-of-plane fiber orientation angles *α* from 3D-sSAXS (***Figure 6A,B***) and the peak distances Δ from SLI (***Figure 6D,E***), both for one vervet brain section (A,D) and one human brain section (B,E). The 3D-arrows in ***Figure 6A*** indicate the 3D orientation of the nerve fibers computed by 3D-sSAXS for four selected regions. The images in ***Figures 6C,F*** show the corresponding 3D fiber orientations from the dMRI measurement of the human brain sample for reference.

The out-of-plane inclination angles from dMRI (***Figure 6C***) highly agree with those obtained from 3D-sSAXS (***Figure 6B***). In both coronal brain sections (vervet and human), the fibers in the corpus callosum (cc) are mostly oriented in-plane (dark blue: *a* < 20°), while fibers in the cingulum (cg) are mostly oriented out-of-plane (light green/yellow: *a >* 40°). Fibers in the vervet fornix (***Figure 6A***) show mostly intermediate inclination angles (light blue: 20° < *a* < 40°).

When comparing the inclination angles to the corresponding SLI peak distances in ***Figure 6D-E*** (evaluated for regions with a single detected fiber orientation), it becomes apparent that regions with inplane fibers (cc) contain many image pixels with large peak distances (blue: Δ > 170°), whereas regions with out-of-plane fibers (cg) contain many image pixels with notably smaller peak distances (green/yellow: Δ < 140°) – especially in the human cingulum. To quantify this effect, we plotted the SLI peak distances against the corresponding 3D-sSAXS inclinations for all evaluated image pixels (see scatter plots in ***Figure 6G**;* data points are shown in similar colors as the corresponding outlines in ***6D-E**;* the insets show the representative SLI azimuthal profiles and corresponding peak distances alongside the dashed-line theoretical prediction). The scatter plots confirm a decreasing peak distance with increasing fiber inclination for most regions, matching the prediction by simulations. The broadly distributed points from the cingulum might be due to the fact that the peak distance in regions with highly inclined fibers is harder to determine due to less pronounced peaks (cf. ***Figure 2C**(iv)*). The data points in the white matter of the human cingulate gyrus (CiG) differ the most from the theoretically predicted curve (brown data points in ***Figure 6G**):* While SAXS yields similarly high fiber inclinations as in the cingulum (magenta data points), the SLI peak distances are much larger (mostly between 160-180°). The large number of gray pixels (surrounded by brown outline in ***Figure 6E***) indicates the existence of crossing fibers. The dMRI orientation distribution functions (***Figure 6F***) reveal indeed that – in addition to the cingulum bundle with highly inclined fibers (in blue) – the cingulate gyrus is interspersed with a transverse rather in-plane fiber bundle (in red), which explains the large SLI peak distances in some regions of the white matter cingulate gyrus.

## Discussion

We performed Scattered Light Imaging (SLI) and small-angle X-ray scattering (SAXS) measurements on the same vervet monkey and human brain sections and compared our human section results to high-resolution *ex vivo* dMRI measurements of the same sample. This allowed us to cross-validate the techniques across different scales and to identify possible limitations - both on the macroscopic scale (dMRI) and the microscopic scale (SLI): Using the 2D-fiber orientations from SLI as high-resolution reference, we found that SAXS yields reliable nerve fiber crossings, while dMRI tends to overestimate the amount of crossing fibers. Taking the main out-of-plane fiber orientations from dMRI and SAXS into account, we could show that SLI provides information about 3D-fiber orientations, but is still limited in the quantification of the out-of-plane angles, especially in regions with crossing fibers. Thus, the combination of the unbiased resolving power of SAXS with the high-resolution power of SLI may best provide a reliable reference for neuronal connectivity maps, and a gold standard to which techniques such as dMRI can be compared.

### Existing methods to identify fiber orientations and tracts

A large variety of neuroimaging techniques exists to study nerve fiber architectures in the post-mortem brain. Some techniques (just as SLI and 2D-SAXS) analyze thin tissue sections to assess brain tissue structures. Histological staining allows to study nerve fiber organizations with fine structural detail (***Amunts et al., 2013; Carriel et al., 2017***), but has limitations in white matter regions with densely packed nerve fibers. Structure-tensor analysis of Nissl-stained histology slides can reveal glial cell orientation along axons (***Schurr 2022***), but the resulting fiber orientations are in 2D; also, dehydration during staining can lead to tissue deformation. Serial electron microscopy (***Eberle & Zeidler, 2018**; **Salo et al., 2021***) enables the analysis of brain tissue structures at highest detail, but is only feasible for very small sample sizes and also requires a complex and specific sample preparation, preventing the reuse of samples for other purposes.

To assess microscopic fiber structures in 3D volumes (without sectioning), tissue clearing followed by labeling and fluorescence microscopy imaging is commonly used. In recent years, it has served as validation for dMRI data (***Marowski et al., 2018; Stolp et al., 2018; Goubran et al., 2019; Leuze et al., 2021***). However, the clearing process causes tissue deformation (***Leuze et al., 2017***). Moreover, it is only feasible for smaller sample sizes (clearing solution and many antibodies cannot fully penetrate large brain samples), and it fails to disentangle densely packed nerve fibers. Other methods to study nerve fiber structures in 3D and microscopic detail (without clearing) are two-photon fluorescence microscopy (***Zong et al., 2017; Costantini et al., 2021***), or optical coherence tomography (***Magnain et al., 2015; Men et al., 2016***) which relies on the back-scattering of light from a tissue block and images the superficial tissue layer before sectioning.

All previously mentioned methods require a directional analysis of the microscopic image data to extract orientation information (structure tensor analysis – ***Khan et al., 2015; Wang et al., 2015***). For this purpose, a kernel including several neighboring image pixels is used, which limits the resolution; also in regions with densely packed nerve fibers intensity gradients are low, limiting analysis.

To directly obtain the axonal orientations, optical coherence tomography can be combined with polarized light (PS-OCT), which exploits the optical anisotropy (birefringence) of myelinated axons to determine their orientations (***Wang et al., 2018; Jones et al., 2020; Jones et al., 2021***). A similar principle is used in polarization microscopy where polarized light is passed through thin brain sections and alterations in the polarization state are measured – a technique known for more than a century (***Brodman, 1903; Fraher et al., 1970***). Recent advances realized polarization microscopy also in reflection mode (***Takata et al., 2018**);* but just as PS-OCT, the techniques only derive 2D fiber orientations. In contrast, threedimensional polarized light imaging determines the 3D-orientations of the nerve fibers (***Axer et al., 2011a; Axer et al., 2011b; Menzel et al., 2015; Zeineh et al., 2017; Stacho et al., 2020; Takemura et al., 2020***) using an advanced signal analysis (***Menzel et al., 2022***) or a tiltable specimen stage (***Schmitz et al., 2018***). However, in contrast to SLI, the techniques yield only a single fiber orientation for each measured tissue voxel, and voxels with multiple crossing fibers yield erroneous fiber orientations (***Dohmen et al, 2015***), while retrieving the out-of-plane angle is also challenging.

All described techniques require subsequent tractography to follow the course of fiber tracts. Tracer studies allow visualization of fiber tracts from their beginning to the end (***Lanciego et al., 2000***), but can only identify specific fiber pathways per experiment and are limited to animal brains. The only ways to follow fiber bundles in *ex vivo* human brains are Klinger’s dissection (***Wysiadecki et al., 2019; Dziedzic et al, 2021***), where accuracy is limited to the macroscopic scale, or tracer injection, which is slow and impractical (***Hevner & Kinney, 1996; Lim et al., 1997a&b***).

### Validation studies of dMRI fiber orientations

To obtain reliable connectivity maps from dMRI, a correct interpretation of the measured diffusion parameters is needed. In recent years, multiple efforts have been undertaken to enhance the interpretation of *in vivo* dMRI data by using post-mortem techniques as validation that provide connectional anatomy maps (***Yendiki et al., 2022***). Techniques used for validation range from histology (***Budde et al., 2012; Seehaus et al., 2015; Schilling et al., 2018***), serial block-face scanning electron microscopy (***Raimo et al., 2018; Salo et al., 2021***), and microscopy of cleared tissue (***Marowski et al., 2018; Goubran et al., 2019; Leuze et al., 2021***) to polarization-sensitive optical coherence tomography (***Wang et al., 2014b; Jones et al., 2020, 2021***) and polarized light imaging (***Caspers et al., 2015; Mollink et al., 2017; Henssen et al., 2019; Caspers & Axer, 2019***).

Multiple studies on simulated data and tracer studies reveal that dMRI tractography often yields false-positive fiber tracts (***Maier-Hein et al., 2017; Schilling et al., 2019; Maffei et al., 2022***). Several studies indicate that dMRI orientations differ up to 20° for secondary fiber orientations and that fiber crossings at angles smaller than 60° cannot be resolved (***Schilling et al., 2016; Schilling et al., 2018***).

SAXS and SLI have both shown the potential to determine secondary (crossing) fiber orientations with a higher precision and smaller crossing angles. As they provide directly structural information across extended fields of view on the same tissue sample, they can serve as a standard validation tool for dMRI-derived fiber orientations, enabling comparisons in different anatomical regions.

### Comparison of SAXS and SLI

Although SAXS and SLI both exploit the scattering of photons to study tissue structures, there exist fundamental differences between them. First, regarding measurement principles (cf. ***Figure 1C,D***): SAXS requires synchrotron radiation and raster-scanning of the sample with the resolution being determined by the beam diameter and scanning step size. SLI can be performed with a simple, inexpensive setup (consisting of an LED display and camera) and provides orientation information for each camera pixel, i.e. with micrometer resolution.

While SAXS uses X-rays with ~0.1nm wavelength interacting with the layered structure of the nerve-surrounding myelin sheath, SLI uses visible light with ~0.5μm wavelength interacting with the directional arrangement of nerve fibers. SLI requires several fibers on top of each other to achieve sufficient signal, whereas SAXS works already on individual (myelinated) fibers (***Inouye et al, 2014***). Also, X-ray scattering always occurs perpendicular to the nerve fibers and the pattern is center-symmetric (cf. ***Figure 2***), while SLI azimuthal profiles with an odd number of peaks cannot be interpreted without taking information from neighboring pixels into account. SAXS allows measurements of samples irrespective of sample thickness, can yield accurate fiber orientations in 3D, and can also be applied tomographically in bulk samples (***Georgiadis et al., 2021***). SLI on the other hand yields much higher in-plane resolutions (here: 33x) without the time-consuming raster-scanning and can be performed with relatively inexpensive equipment in a standard laboratory.

Despite these differences, SAXS and SLI also have much in common. They are both orientation-specific methods: they directly probe the fiber orientation, without an intermediate step of imaging the tissue structures as in optical or electron microscopy, or using a proxy such as anisotropic water diffusivity in dMRI. This enables to reliably determine the nerve fiber orientations also in regions with densely packed, multi-directional fibers. They also result in similar azimuthal profiles for in-plane fibers, and, as here, the same software can be used to determine peak positions for both techniques. At the same time, both techniques can image similarly-prepared tissue sections, without any staining or labeling, and they are nondestructive, enabling sample reuse.

### Identification of false-positive fiber tracts in dMRI

The 2D fiber orientations from the highly-specific SAXS measurement corresponded very well to those from the high-resolution SLI measurement (***Figure 3-4***), demonstrating the ability of both techniques to serve as ground truth for in-plane fiber orientations in complex brain tissue structures. Registering dMRI, 3D-sSAXS, and SLI data sets of a human brain sample enabled comparisons of fiber orientations from all three methods (***Figure 5*** and ***Figure 5–figure supplements 1-3***). When comparing the 3D-sSAXS fiber orientations of two brain sections with the corresponding dMRI fiber orientation distributions of the entire tissue sample, we observed a very high correspondence between the primary fiber orientations for each voxel: the dot product is highly skewed towards one, denoting almost perfect co-alignment (***Figure 5D,E*** and ***Figure 5– figure supplement 2***), similar to what had been shown previously in mouse brain (***Georgiadis et al., 2020***). The regions with low dot product (colored in blue in ***Figure 5E*** and ***Figure 5–figure supplement 2***) are regions with two strong crossing fiber populations (cf. ***Figure 5G***), so correspondence of primary orientation is expectedly low. When considering crossing fibers, in the most challenging regions of the corona radiata, the fiber orientations from dMRI and SAXS also seemed to be in high agreement (***Figure 5– figure supplement 3B***).

However, we observed a discrepancy in regions with more homogeneously distributed fibers, such as in the corpus callosum (arrows in ***Figure 5G**):* Diffusion MRI seemed to consistently yield a secondary fiber population perpendicular to the main one, albeit with much smaller magnitude (***Figure 5F*** (ii) and (iii)). X-ray scattering did not show such a crossing (***Figure 5G*** right), which was also missing in the micron-resolution scattered light imaging (***Figure 5H***), both showing unidirectional fibers in these voxels (regions 1 and 2). Two effects might explain the sensitivity to possibly non-existent fiber populations in the dMRI data sets: First, there is a known issue of possibly false-positive/spurious fiber populations due to the overfitting of the response function to diffusion data (***Guo et al., 2021; Baete et al., 2019***), especially in the presence of noise. This is also visible in the diffusion MRI fiber orientations distributions in ***Mollink et al. (2017)***, Figure 8, which are not present in PLI or histology. Second, upon looking closely at the micrometer-resolution microscopy images in the corpus callosum region, small vessels running perpendicular to the fiber orientations can be observed (yellow arrows in ***Figure 5H***). It is possible that the small, aligned structures affect the diffusion MRI signal, with some population of water molecules being constrained to diffuse in the direction of these vessel walls. Even a small such effect could potentially give rise to small artificial fiber populations in these directions. A possible solution would be to increase the threshold of secondary lobes prior to running tractography algorithms, as suggested in ***Maffei et al. (2022***). However, this approach, while increasing the specificity, might decrease the sensitivity for the cases where there exist actual but less prominent secondary fiber populations.

Such phenomena stress the need for approaches that use micro-structural models to decouple the contributions from intra- and extra-cellular water (***Jelescu and Budde, 2017***). Using such models could help to separate the hindered diffusion close to these vessel walls and the restricted diffusion within the axons, making the dMRI-derived fiber orientations insensitive to such signals and thus more axon-specific. Selection of the optimum model that best eliminates these contributions is not within the scope of the current manuscript, but our results show that research in this direction should be pursued in the future, using the directly structural, fiber-specific and/or micrometer-resolution methods presented here as ground-truth data to refine the models.

### Experimental validation of out-of-plane fibers in SLI

With the combined measurement of SLI and 3D-sSAXS (and dMRI for the human sample), we were able to provide experimental validation of the predicted decrease in SLI peak distance with increasing fiber inclination. However, it also became apparent that the quantification of fiber inclination based on SLI peak distance alone is challenging: while regions with steep fibers (inclinations > 70°) can be clearly identified by a high degree of scattering and small peak distances (< 90°), the moderate decrease in peak distance for fibers with up to 60° inclination together with the large distribution of measured values (cf. ***Figure 6G***) makes a clear assignment between peak distance and inclination practically impossible. Our study suggests that SLI also has limitations when it comes to regions with inclined crossing fibers (cf. ***Figure 6E-F***, and ***G*** on the right). To improve the interpretation, more advanced algorithms are needed. Machine learning models, trained on simulated data sets, could help to improve the interpretation of measured scattering patterns from SLI and yield more reliable estimates (suggested by ***Vaca et al., 2022***).

### Conclusion

Disentangling the highly complex nerve fiber architecture of the brain requires a combination of dedicated, multi-scale imaging techniques. We here provide a framework that enables combined measurements of scattered light and X-ray scattering (SLI and SAXS) on the same brain tissue sample, with high agreement between the two methods. The high-resolution properties of the former combined with the high-specificity of the latter enables the detailed reconstruction of multiple nerve fiber orientations for each image pixel, which can provide providing unprecedented insights into brain circuitry. The unique cross-validation of SLI, SAXS, and diffusion MRI on the same tissue sample revealed high agreement between the methods, but also false-positive crossings in MRI. Furthermore, it allowed the experimental validation of out-of-plane fiber orientations in SLI, paving the way for a more detailed reconstruction of 3D nerve fiber pathways in the brain. Due to the simple setup of SLI, any SAXS measurement of a tissue section can easily be combined with a corresponding SLI measurement, significantly enhancing the reconstruction of nerve fiber pathways in the brain, especially in regions with complex fiber crossings.

## Materials and methods

### Vervet brain sample preparation

The vervet monkey brain was obtained from a healthy 2.4-year-old adult male in accordance with the Wake Forest Institutional Animal Care and Use Committee (IACUC #A11-219). Euthanasia procedures conformed to the AVMA Guidelines for the Euthanasia of Animals. All animal procedures were in accordance with the National Institutes of Health guidelines for the use and care of laboratory animals and in compliance with the ARRIVE guidelines. The brain was removed from the skull within 24 hours after death, 4% formaldehyde-fixed for several weeks, cryo-protected in 20% glycerin and 2% dimethyl sulfoxide, deeply frozen, and coronally cut from the front to the back into 60μm-thick sections using a cryostat microtome (*Polycut CM 3500*, Leica Microsystems, Germany). The brain sections were mounted on glass slides, embedded in 20% glycerin, and cover-slipped. Two sections from the middle (no. 511 and 501) were selected for further evaluation (see ***Figure 2A*** and ***Figure 2–supplement figure 1A***). A region from the right hemisphere (16.4×10.9mm^2^) - containing part of the corona radiata, corpus callosum, cingulum and fornix – was measured with SLI several months afterwards (cf. ***Figure 2B*** and ***Figure 2–supplement figure 1B***). For 3D-sSAXS, the brain sections were removed from the glass slides, re-immersed in phosphate-buffered solution (PBS) for two weeks, placed in-between two 170μm-thick (#1.5) cover slips, sealed, and measured in a comparable region (19.0×10.9mm^2^, cf. ***Figure 2C*** and ***Figure 2–supplement figure 1C***).

### Human brain sample preparation

The human brain (66 year-old female with no known neurological disorders) was obtained from the Stanford ADRC Biobank, which follows procedures of the Stanford Medicine IRB-approved protocol #33727, including a written informed brain donation consent of the subject or their next of kin or legal representative. The brain was removed from the skull within 24 hours, fixed for 19 days in 4% formaledhyde (10% neutral buffered formalin), coronally cut into 1 cm-thick slabs, and stored in PBS for five years. From the left hemisphere, a 3.5×3.5×1cm^3^ specimen – containing part of the corona radiata, corpus callosum, and cingulum – was excised (cf. ***Figure 5A***). For dMRI, the specimen was degassed and scanned in fomblin. Five weeks later, the anterior and posterior part of the tissue was cut with a vibratome (VT1000S, Leica Microsystems, Germany) into 80μm-thick sections. Two sections (no. 18 from the posterior side and no. 20 from the anterior side) were selected for further evaluation. For 3D-sSAXS, the brain sections were placed in-between two 150μm-thick (#1) cover slips and measured in a center region of 28.0×18.9mm^2^ for no. 18 (red rectangle in ***Figure 5B***) and 28.0×20.1mm^2^ for no. 20. For SLI, the brain sections were removed from inbetween the cover slips, mounted on glass slides with 20% glycerin, cover-slipped, and measured ten weeks afterwards in a region of 16.4×10.9 mm^2^ containing corpus callosum and cingulum (cf. blue rectangle in ***Figure 5B***).

### Scattered Light Imaging

The SLI measurements (cf. ***Figure 1D***) were performed using an LED display (*Absen Polaris 3.9pro In/Outdoor LED Cabinet*, Shenzen Absen Optoelectronic Co., Ltd., China) with 128×128 individually controllable RGB-LEDs with a pixel pitch of 3.9mm and a sustained brightness of 5000cd/m^2^ as light source. The images were recorded with a CCD camera (*BASLER acA5472-17uc*, Basler AG, Germany) with 5472×3648 pixels and an objective lens (*Rodenstock Apo-Rodagon-D120*, Rodenstock GmbH, Germany) with 120mm focal length and 24.3cm full working distance, yielding an in-plane resolution of 3.0μm/px and a field of view of 16.4×10.9 mm^2^. The distance between light source and sample was set to approximately 16cm, the distance between sample and camera to approximately 50cm.

The SLI scatterometry measurement (used to generate the scattering patterns in upper ***Figure 2C***) was performed as described in ***Menzel et al. (2021b***: A square of 2×2 illuminated RGB-LEDs (white light) was moved over the LED display in 1-LED steps for a square grid of 80×80 different positions, and an image was taken for every position of the square with an exposure time of 1sec. For each position of an illuminating square of LEDs, four shots were recorded and averaged to reduce noise. In the end, for each point of the sample a scattering pattern with 80×80 pixels was assembled (cf. ***Figure 1D*** on the right): The upper left pixel in the scattering pattern shows the intensity of the selected point in the image that was recorded when illuminating the sample from the upper left corner of the display, and so on. The azimuthal profiles in upper ***Figure 2C*** were generated by integrating the values of the scattering pattern from the center (point of maximum intensity) to the outer border of the pattern and plotting the resulting value *I*(φ) against the respective azimuthal angle (φ=0°,1°, … 359°).

The angular SLI measurements (used to generate the SLI parameter maps in ***Figures 3-6, Figure 2–figure supplement 1*** and ***Figure 3–figure supplement 1***) were performed as described in ***Menzel et al. (2021a)***: A rectangle of illuminated green LEDs (2.4×4 cm^2^) was moved along a circle with a fixed polar angle of illumination (*θ*=45°) and steps of Δφ=15°. For every position of the rectangle (φ=0°, 15°, … 345°), an image was taken with an exposure time of 0.5sec. The resulting series of 24 images (containing azimuthal profiles, i.e. intensity values for each measured azimuthal angle φ for each image pixel) was processed with the software *SLIX* (Scattered Light Imaging ToolboX) v2.4.0 (https://github.com/3d-pli/SLIX) to generate the orientational parameter maps, as described below.

### 3D-scanning small-angle X-ray scattering

3D-sSAXS (***Georgiadis et al., 2015**; **Georgiadis et al., 2020***) was performed at beamline 4-2 of the Stanford Synchrotron Radiation Lightsource, SLAC National Accelerator Laboratory, with a beam of photon energy Epiroto⊓=15keV. The vervet brain sections were measured (cf. ***Figure 1C***) with a beam diameter of 100μm, an exposure time of 0.7sec, rotation angles θ = [0°, +/-15°, … +/-60°], and a field of view of 19.0×10.9mm^2^ at 100μm x- and y-steps. The human brain sections were measured with a beam diameter of 150μm, an exposure time of 0.4sec, rotation angles θ = [0°, +/-10°, … +/-70°], and a field of view of 28.0×18.9mm^2^ (anterior section no. 18) and 28.0×20.1mm^2^ (posterior section no. 20) at 150μm x- and y-steps.

To compute the in-plane fiber orientations (shown in ***Figures 3–5**, **Figure 3–figure supplement 1*** and ***Figure5–figure supplement 3***), azimuthal profiles were generated for each scattering pattern of the θ=0°- measurement (cf. lower ***Figure 2C***) and analyzed by the same SLIX software, as described below. To generate the azimuthal profiles, the scattering patterns were divided into Δφ=5°-segments, the intensity values were summed for each segment, and the resulting values were plotted against the corresponding average φ-value. The known center-symmetry of the SAXS scattering patterns was exploited to account for missing parts due to detector electronics.

The out-of-plane fiber inclination angles (***Figures 5C*** and ***6A-B***) were computed by analyzing the scattering patterns obtained from 3D-sSAXS measurements at different sample rotation angles, as described in ***Georgiadis et al. (2020***).

### Diffusion magnetic resonance imaging

The dMRI measurement was performed on a Bruker 11.7 T scanner, using a 12-segment spin-echo echo planar imaging (SE-EPI) sequence at 200μm isotropic voxels, repetition time TR=400ms, echo time TE=40ms, diffusion separation time δ=7ms, diffusion time Δ=40ms, field of view FOV=40×36×21 mm^3^, at 200 diffusion-weighted *q*-space points (20@b=1ms/μm^2^, 40@b=2ms/μm^2^, 60@b=5ms/μm^2^, 80@b=10ms/μm^2^) and 20@b=0ms/μm^2^. First, data were denoised and corrected for Gibbs artifacts (***Ades-Aron et al., 2018**; **Veraart et al., 2016***). Then, volumes were b-value-averaged, and registered to the initial b0 volume using *FSL FLIRT* (https://fsl.fmrib.ox.ac.uk/fsl/fslwiki/FLIRT; ***Jenkinson et al., 2002***) with mutual information as cost function and a spline interpolation. After registration to the SLI and SAXS (see corresponding ‘Image registration’ Methods section), fiber responses and orientation distributions were computed using the *dwi2response* and *dwi2fod* functions in MRtrix3 (https://www.mrtrix.org/) –employing *dhollander* and *msmt_csd* (multi-tissue, multi-shell constrained spherical deconvolution) algorithms respectively– and visualized in *mrview*. The dMRI-derived output fiber orientation distributions for each voxel were sampled at the plane of the vibratome section in 5°-steps using MRtrix3’s *sh2amp* command, which was then used as input to the SLIX software package for computing in-plane fiber orientations including crossings. For main fiber orientations, diffusion tensor imaging (DTI) processing was performed using FSL’s *DTIFIT* function (https://fsl.fmrib.ox.ac.uk/fsl/fslwiki/FDT/UserGuide#DTIFIT). For the dMRI parametric maps, the DESIGNER pipeline (https://github.com/NYU-DiffusionMRI/DESIGNER; ***Ades-Aron et al., 2018***) was used to compute diffusivity, kurtosis and white matter tract integrity parameters (***Fieremans et al., 2011***).

### Generation of orientational parameter maps

The azimuthal profiles from angular SLI, 3D-sSAXS, and dMRI were processed with SLIX in order to generate various parameter maps (***Figure 2–figure supplement 1***) and to determine the in-plane nerve fiber orientations. The analysis of the profiles and the software are described in ***Menzel et al. (2021a)*** and ***Reuter and Menzel** (**2020***) in more detail. The software determines the positions of the peaks for each image pixel (azimuthal profile). The peak prominence (***Figure 2–figure supplement 1***, 4^th^ row) was determined as the vertical distance between the top of the peak and the higher of the two neighboring minima. Only peaks with a prominence larger than 8% of the total signal amplitude (max – min) were considered for evaluation.

The peak width (***Figure 2–figure supplement 1***, last row) was computed as the full width of the peak at a height corresponding to the peak height minus half of the peak prominence. The in-plane fiber orientation *φ (**Figures 3-5**, **Figure 3–figure supplement 1*** and ***Figure5–figure supplement 3***) was computed as the midposition between peaks that lie 180° +/- 35° apart. To better analyze multiple crossing fiber orientations, the in-plane fiber orientations were visualized as colored lines and displayed on top of each other (cf. ***Figure 3*** and ***Figure 3–figure supplement 1***). The peak distance (***Figure 6D-E***) was computed as the distance between two peaks, for profiles with no more than two peaks (profiles with one peak yield zero peak distance).

### Image registration

To register 3D-sSAXS onto SLI (***Figures 4,6***), the 3D-sSAXS parameter maps were upscaled to the SLI pixel size. Linear registration of 3D-sSAXS to SLI sections was performed using FSL FLIRT (https://fsl.fmrib.ox.ac.uk/fsl/fslwiki/FLIRT), while angular information and 3D vectors were rotated accordingly. For registering dMRI onto SAXS (***Figure 5*** and ***Figure 5–figure supplement 2-3***), first the matching plane for each human brain section was identified manually in the scanned MRI volume (different plane for each human brain section), and FSL FLIRT linear registration with 12 degrees of freedom was used for precise alignment of the 2D images. Then, the entire dMRI data set was transformed using the identified rotation and translation parameters (twice, once for each section), and the b-vectors were rotated correspondingly. The MRI sections corresponding to the vibratome section plane were isolated and further analyzed as explained in the ‘Diffusion magnetic resonance imaging’ Methods section.

## Acknowledgements

We thank Laura Pisani from the Stanford Center for Innovation in In vivo Imaging (SCi3) and Kristin Garlund from Bruker BioSpin USA for assistance with the dMRI measurements, Roger Woods from the UCLA Brain Research Institute and Donald Born from Stanford Pathology for providing the vervet and human brain samples, and the laboratory team from the Institute of Neuroscience and Medicine (INM-1), Forschungszentrum Jülich, for the preparation of the vervet and human brain sections for the SLI measurements. This work was supported by the National Institutes of Health (NIH) under award numbers R01NS088040, P41EB017183, R01AG061120-01, R01MH092311, and 5P40OD010965, by the Helmholtz Association port-folio theme “Supercomputing and Modeling for the Human Brain”, by the European Union’s Horizon 2020 Research and Innovation Programme under Grant Agreement No. 945539 (“Human Brain Project” SGA3), and by the Deutsche Forschungsgemeinschaft (DFG, German Research Foundation). The Stanford Synchrotron Radiation Lightsource, SLAC National Accelerator Laboratory, is supported by the U.S. Department of Energy, Office of Science, Office of Basic Energy Sciences under Contract No. DE-AC02-76SF00515. The SSRL Structural Molecular Biology Program is supported by the DOE Office of Biological and Environmental Research, and by the National Institutes of Health, National Institute of General Medical Sciences (P30GM133894). The Pilatus detector at beamline 4-2 at SSRL was funded under National Institutes of Health Grant S10OD021512. M.M. received funding from the Helmholtz Doctoral Prize 2019 and the Klaus Tschira Stiftung gGmbH.

## Additional information

### Competing interests

The authors declare that no competing interests exist.

### Author contributions

**Miriam Menzel**, Conceptualization, Methodology, Formal analysis, Investigation, Visualization, Supervision, Funding acquisition, Writing–original draft, Writing–review and editing; **David Gräßel**, **Ivan Rajkovic**, Investigation, Writing–review and editing; **Michael Zeineh**, Conceptualization, Formal analysis, Supervision, Funding acquisition, Resources, Writing–review and editing; **Marios Georgiadis**, Conceptualization, Methodology, Formal analysis, Investigation, Visualization, Supervision, Funding acquisition, Writing–original draft, Writing–review and editing

## Additional files

### Code and data availability

All software used for image processing is open-source and described in the Methods section (with URL links). Data analysis for Figures 4 and 6 was performed with Fiji (https://fiji.sc/Fiji), for Figure 5 and supplementary figures using Matlab 2021b (Mathworks, USA). All data and code supporting the findings of this study are available from the corresponding authors upon request.

## Figure Supplements

**Figure 2–figure supplement 1.**
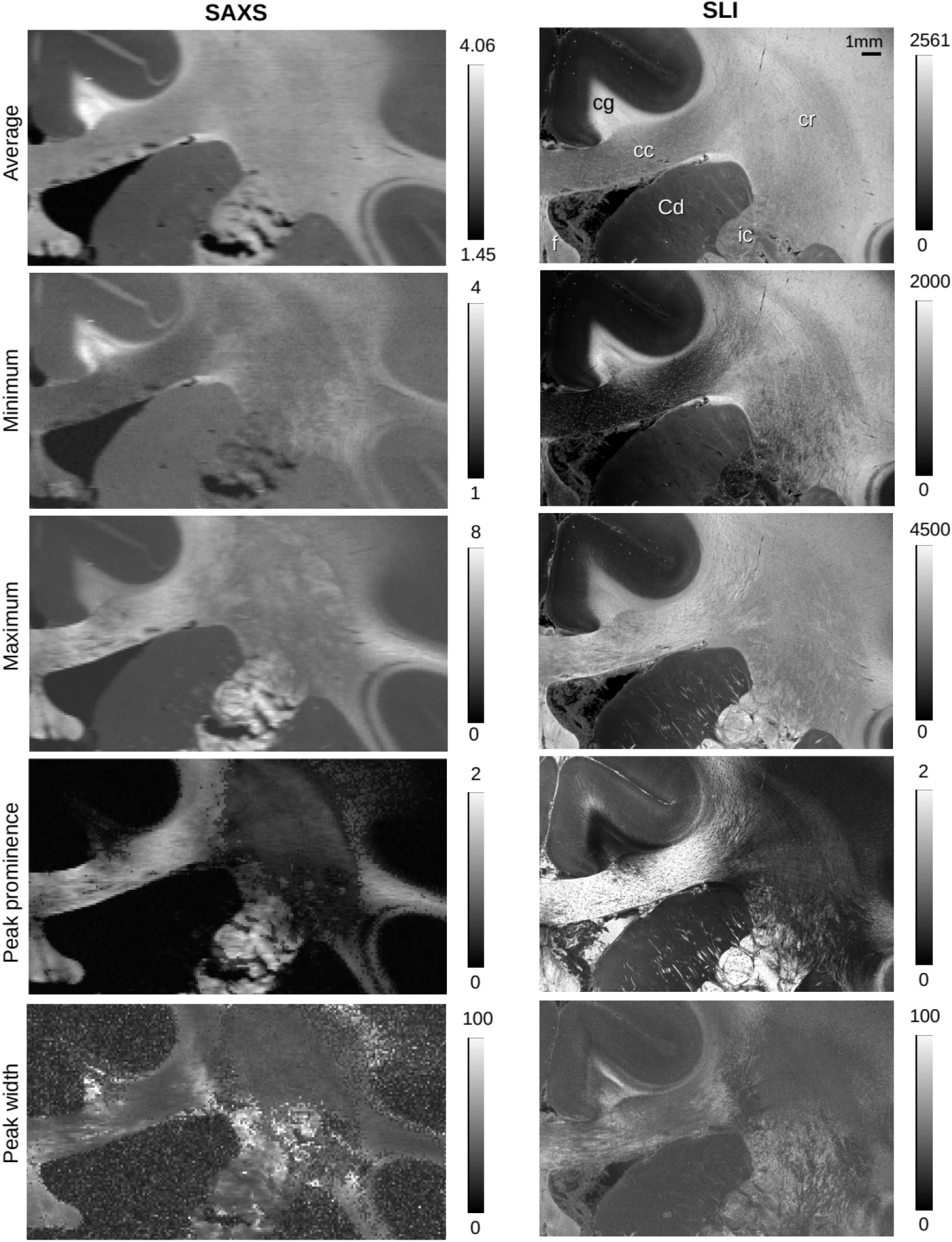
Parameter maps obtained from SAXS and SLI azimuthal profiles for vervet brain section no. 511. The top images show the average, maximum, and minimum values of the azimuthal profiles for each image pixel. The lower images show the mean prominence and width of the peaks in the azimuthal profiles. The images show a similar behavior corresponding to the azimuthal profiles shown in **Figure 2C**: Out-of-plane nerve fibers in the cingulum yield high average scattered light intensities with small signal amplitude (max-min), small peak prominence, and large peak width. In-plane nerve fibers in the corpus callosum yield a large signal amplitude, high peak prominence, and small peak width. In-plane crossing nerve fibers in the corona radiata yield a smaller signal amplitude and less prominent peaks.

**Figure 3–figure supplement 1.**
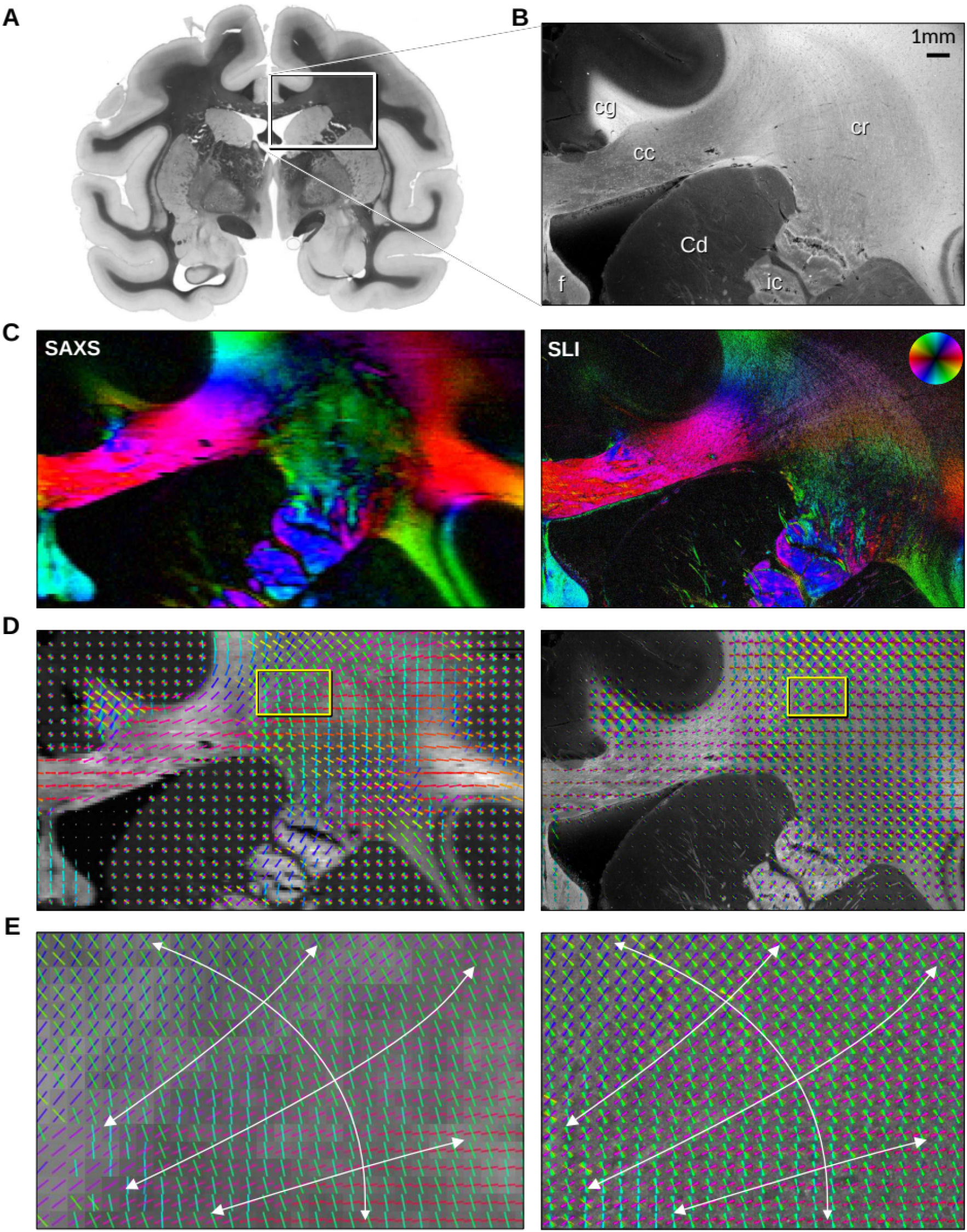
In-plane fiber orientations from SAXS and SLI measurements of vervet brain section no. 501. (**A**) Transmittance image of the whole section. (**B**) Average scattered light intensity of the investigated region (cc: corpus callosum, cr: corona radiata, cg: cingulum, Cd: caudate nucleus, f: fornix, ic: internal capsule). (**C**) Fiber orientation maps showing the predominant fiber orientation for each image pixel in different colors (see color wheel in upper right): px=100μm (SAXS), px=3μm (SLI). (**D**) Fiber orientations displayed as colored lines for 5×5 px (SAXS) and 165×165 px (SLI) superimposed. (**E**) Enlarged region of the corona radiata, showing fiber orientations as colored lines for 1×1 px (SAXS) and 33×33 px (SLI) superimposed. The white arrows indicate the overall course of the fiber vectors.

**Figure 5–figure supplement 1.**
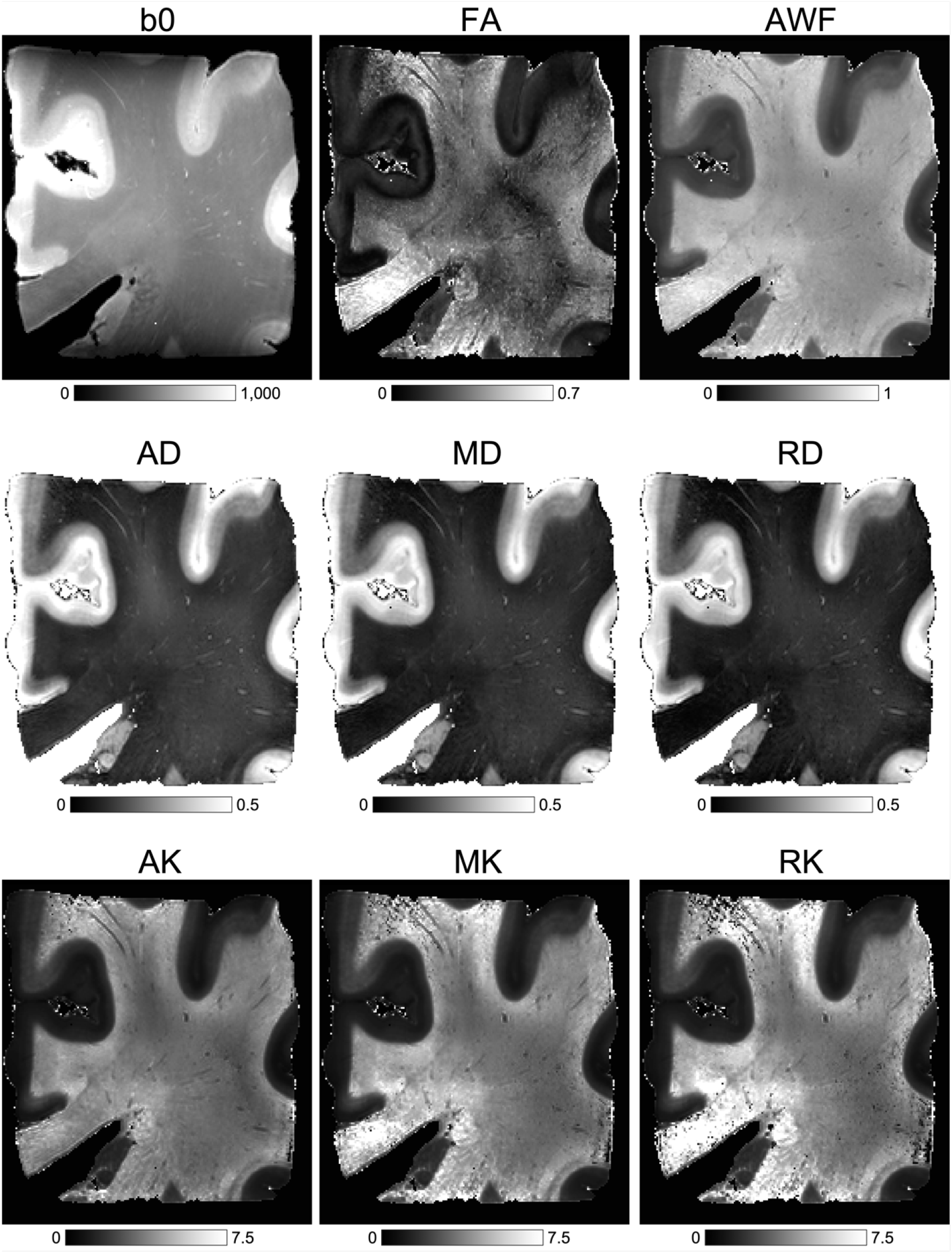
Anatomic (b0 – T2w) and diffusion MRI-based metrics. Calculated using the DESIGNER pipeline, which includes kurtosis and white matter tract integrity (WMTI) metrics. Fractional anisotropy (FA), axonal water fraction (AWF), axial, mean and radial diffusivity (AD, MD, RD), axial, mean, and radial kurtosis (AK, MK, RK).

**Figure 5–figure supplement 2.**
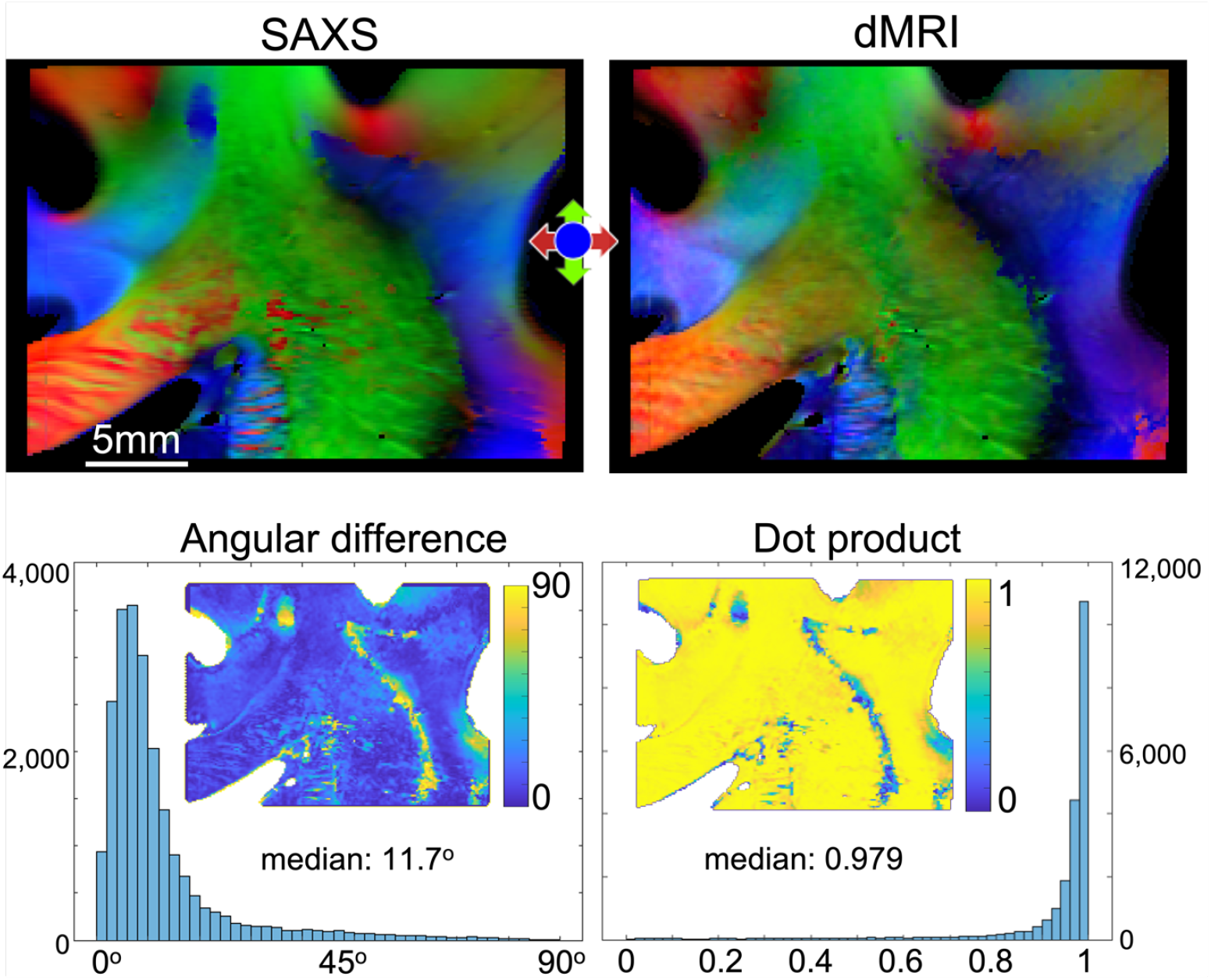
SAXS-dMRI comparison for anterior section (A20). The top part shows the 3-dimensional orientations of the fibers retrieved by 3D-sSAXS and dMRI, respectively. The bottom part quantifies the difference in the angles retrieved by the two methods. To the left, the absolute angular difference is plotted as a histogram and mapped on the section. To the right, the angular difference is quantified in the form of a dot product. The median angular difference found is 11.7°, which corresponds to a dot product of 0.970.

**Figure 5–figure supplement 3.**
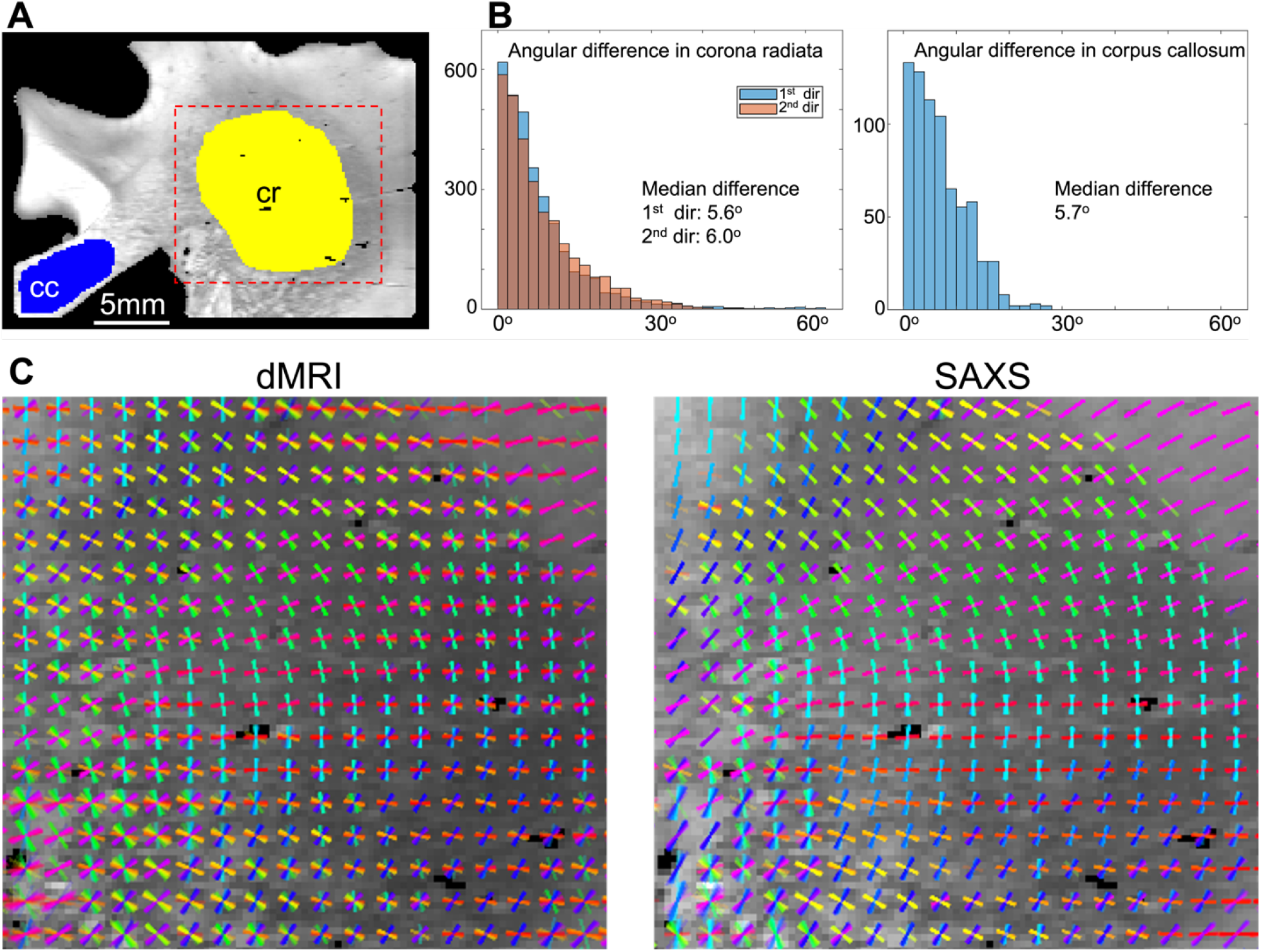
Quantifying in-plane angular differences between SAXS and dMRI for the corpus callosum (cc) and corona radiata (cr) areas of posterior section (B18). (**A**) Map of the scattering intensity of the section, depicting the areas where quantification was performed. (**B**) Left, the histograms of the angular differences of the first and second fiber direction in the corona radiata are overlaid, showing very similar results (5.6° difference for the first direction, 6° for the second). Right, the same quantification for the corpus callosum area, showing a difference of 5.7°. (**C)** Zoom-in to the fiber orientations in the corona radiata, retrieved by dMRI and SAXS (orientations of 5×5 pixels are displayed on top of each other).

## Notes

### Competing Interest Statement

The authors have declared no competing interest.

